# PPIKB: A Comprehensive Knowledge Base and Analysis Platform for Protein–Peptide Interactions Based on Literature and Patents

**DOI:** 10.1101/2025.06.09.658607

**Authors:** Ning Zhu, Yanyu Ming, Chengyun Zhang, Cao Sen, Chongyang Li, Jingjing Guo, Hongliang Duan

## Abstract

Protein–protein interactions (PPIs) underpin a multitude of fundamental biological processes and represent critical therapeutic targets. Peptide compounds—owing to their structural flexibility, high target specificity, and the combined advantages of small molecules and biologics—have emerged as powerful modulators of these interactions. However, the exponential growth of scientific publications and patent filings poses a significant challenge for the systematic integration of protein–peptide interaction (PPepI) data. To address this, we implemented a high-throughput extraction pipeline powered by large language models (LLMs) to harvest affinity data from both research articles and patents, followed by rigorous manual curation. Applying this approach to 4,241 research papers and 483 patents published between 1972 and 2025, we compiled 21,760 manually validated quantitative affinity entries encompassing 2,013 unique protein targets derived from human, mouse, rat, viral, bacterial, and other species. In parallel, we consolidated 18,005 structural entries of protein–peptide complexes from the Protein Data Bank and associated literature. Leveraging these comprehensive datasets, we developed PPIKB, a user-friendly database and analysis platform that enables efficient querying, visualization, and in-depth analysis of protein–peptide interactions. PPIKB also provides a suite of tools tailored to facilitate peptide-based drug discovery, protein engineering, and systems-level biological research. By combining state-of-the-art text-mining technologies with expert validation, PPIKB sets a new standard for accessibility and utility in the field of protein–peptide interaction data resources. Access PPIKB at: https://ppikb.duanlab.ac

## Introduction

Protein–protein interactions (PPIs) lie at the core of virtually all biological processes— including signal transduction, intercellular communication, immune responses, and metabolic regulation—and their dysregulation often drives disease pathogenesis, making PPIs attractive therapeutic targets^1,2^. Traditional small-molecule drugs have a long clinical history owing to their low production costs, oral bioavailability, favorable cell-membrane permeability, and generally excellent safety and patient compliance. Whether derived from natural products or produced synthetically, small molecules remain more cost-effective than peptides or biologics^3–5^; however, their relatively small size, while advantageous for intracellular access, limits their ability to inhibit the large, flat surfaces that typify many PPIs, posing a fundamental challenge for PPI modulation.In contrast, peptide therapeutics offer structural flexibility, high binding specificity, low immunogenicity, and moderate manufacturing costs, making them powerful candidates for targeting PPIs^6–9^. Beyond therapeutics, peptides serve as highly selective recognition elements in biosensors and diagnostic reagents: their excellent biocompatibility, stability, and ease of chemical modification have driven advances in peptide-based electrochemical biosensors, which combine high sensitivity, rapid response, and low cost to enable point-of-care diagnostics and real-time health monitoring^10–13^.

Recent years have witnessed dramatic strides in artificial intelligence (AI) for drug discovery, particularly in protein structure prediction. General-purpose tools such as DeepMind’s AlphaFold and BakerLab’s RoseTTAFold have revolutionized our ability to model protein conformations^14,15^. Building on these foundations, researchers have developed AI models expressly tailored to peptide-specific challenges. Early fragment-based approaches, exemplified by the PEP-FOLD series, focus on monomeric peptide folding^16^; our HighFold pipeline further refines cyclic-peptide and peptide–protein complex modeling, furnishing critical structural insights for mechanism elucidation and lead optimization^17^. Models such as AFsample2^18^ address peptide flexibility by predicting multiple conformers, deepening our understanding of dynamic behavior.

AI has also transformed de novo design and optimization of functional peptides. Diffusion-based generative frameworks (e.g., RFdiffusion) can craft novel protein scaffolds and peptide sequences targeting specific binding motifs^19^. Tools like PepINVENT explore expansive peptide chemical spaces—including noncanonical amino acids—to achieve predefined objectives such as enhanced affinity, stability, or solubility^20^. Integrating generative models (e.g., VAE- or GRU-based architectures) with molecular docking and molecular dynamics (e.g., RosettaFlexPepDock) has led to experimentally validated, high-affinity inhibitors against targets such as β-catenin^21,22^. Reinforcement-learning strategies—such as HighPlay^23^, which combines HighFold structure prediction with sequence optimization—enable simultaneous refinement of peptide sequence and binding characteristics. Meanwhile, transformer-based predictors (e.g., TPepPro), which fuse sequence and structural features, have markedly improved the accuracy of protein–peptide interaction forecasts, reducing experimental screening burdens and enhancing hit rates^24^.

Despite these methodological advances, the development, benchmarking, and validation of AI-driven models—and the broader pipeline for peptide discovery— depend critically on access to high-quality, large-scale protein–peptide interaction datasets that integrate sequence data, precise quantitative affinity measurements, and high-resolution structural information. Constructing and continuously expanding such a comprehensive knowledge base is therefore essential for overcoming current R&D bottlenecks, expediting the discovery of peptide-based PPI modulators, and driving iterative improvements in AI predictive and design tools. However, the explosive growth of publications and patents renders manual curation increasingly impractical, underscoring an urgent need for automated, AI-assisted data-extraction technologies.

Large language models (LLMs) such as GPT-3, GPT-4, LLaMA, and PaLM have recently demonstrated exceptional capabilities in extracting structured data from unstructured text by recognizing named entities and their relationships^25^. For example, joint fine-tuning of GPT-3 and LLaMA-2 has enabled efficient multi-tiered data extraction in materials chemistry, while LLaMA-driven pipelines have achieved near-human accuracy in extracting pathological features from breast cancer reports^26^. Interactive platforms like SciDaSynth facilitate user-guided, table-based knowledge synthesis^27^, and frameworks such as LLMs4SchemaDiscovery automate domain ontology generation through human–model collaboration. Across chemistry, materials science, and biomedicine^28^, LLM-based approaches have proven their power to convert vast literature corpora into high-value structured datasets^29,30^.

Existing PPI and peptide resources fall into two main categories. Structure-only databases derived from the Protein Data Bank^31^—including PepBind^32^ (≈3,100 entries; now inaccessible), Propedia V2.3^33^ (>49,300 clustered complexes), and PepBDB (13299 complexes with computed interface metrics)—lack comprehensive affinity or patent data^34^. Binding-affinity repositories such as PDBbind^35^ and BindingDB^36^ provide quantitative data for protein–ligand and protein–small-molecule interactions but systematically overlook peptide ligands and, in some cases, are proprietary. Consequently, none of these resources fully meet the community’s need for an integrated, accurate, and scalable protein–peptide interaction knowledge base.

To address these limitations, we have developed PPIKB: a comprehensive knowledge base and analysis platform for protein–peptide interactions that combines high-throughput LLM-based extraction from literature and patents with rigorous expert validation. PPIKB encompasses two principal datasets: (1) quantitative affinity data— 21,760 manually validated entries (3,142 with corresponding crystal structures) extracted from 4,241 research articles and 483 patents published from 1972 to the present, covering 2,013 unique protein targets across diverse species; and (2) structural data—18,005 PDB-derived protein–peptide complex entries with detailed interaction analyses computed via PLIP. Accessible at https://ppikb.duanlab.ac, PPIKB features an intuitive web interface supporting text and structure search, classification browsing, rich visualization, bulk data download, and integrated analysis tools (Foldseek^37^, BLAST^38^, SMILES similarity). By uniting advanced text-mining methodologies with stringent manual curation, PPIKB establishes a new benchmark in data breadth, depth, and usability—providing a unified, high-quality platform to accelerate peptide drug discovery, computational modeling, and fundamental biological research.

## Methods

### 1. Data Sources and Selection Criteria

To ensure the comprehensiveness and systematicity of PPIKB’s data collection, we queried multiple leading scientific literature databases—PubMed, Google Scholar, and Scopus—and retrieved patent information from lens.org, Google Patents, and the patent offices of major jurisdictions (United States, China, Japan, and the European Union). Our search strategy centered on the core concepts of protein–peptide interactions and binding affinity, employing terms such as “protein peptide interaction,” “affinity,” “surface plasmon resonance,” “biolayer interferometry,” “fluorescence polarization,” “enzyme activity assay,” “microscale thermophoresis,” and quantitative descriptors “Kd,” “Ki,” and “IC50.” Using these targeted queries, we identified 4,241 research articles reporting affinity measurements and 483 related patents. In parallel, we systematically downloaded all publicly available three-dimensional protein–peptide complex structures from the Protein Data Bank^31^.

PPIKB’s inclusion criteria were as follows:

1. Peptide sequence clarity: the peptide sequence must be explicitly specified and contain no more than 100 amino acids.
2. Experimental affinity data: reported affinity values must be numerical and derived from wet-lab experiments (e.g., Kd = 1.5 nM).
3. Protein target definition: the protein target must have a clearly defined sequence, either directly provided or unambiguously inferred from gene and species information in the source publication or patent.
4. Source verification: all affinity data must originate from peer-reviewed articles or granted patents.

To further enrich each database entry, we computed key physicochemical properties of peptides using bioinformatics tools such as RDKit^39^ and peptides. Protein–peptide interface interactions were predicted with PLIP^40^, and protein–protein network data were integrated from the STRING database^41^. These stringent selection and annotation workflows collectively guarantee the high quality, reliability, and informational depth of PPIKB, furnishing researchers with a rich and trustworthy resource for protein– peptide interaction data.

### 2. LLM-Based Data Extraction

We evaluated the feasibility and accuracy of large language models (LLMs) for automated extraction of protein–ligand affinity data by conducting a head-to-head comparison of state-of-the-art models. Our focus was on extracting affinity values embedded within deeply unstructured scientific text—an area where both manual curation and traditional automated pipelines face significant challenges. We used Affinity Benchmark v5.5 as the reference dataset and randomly selected 150 articles containing experimentally measured affinity data.

The LLMs assessed were O3-mini-2025-01-31, Claude-3-5-haiku, Deepseek-reasoner, and Gemini-1.5-pro, each exemplifying cutting-edge natural language processing capabilities. Since these models exhibit limitations when ingesting native PDF files, we first converted all source PDFs into plain text via the PyMuPDF library, preserving page order and essential layout information while stripping reference sections. This preprocessing step maximized retention of critical contextual cues and minimized unnecessary token consumption and computational overhead, thereby establishing a fair and efficient foundation for model evaluation.

As illustrated in Figure 1, we devised an identical prompt for each model to extract, in tab-separated-value (TSV) format, the ligand name, receptor protein name, receptor species, affinity value, experimental method, and associated PDB ID—one record per line. To suppress output variability and ensure reproducibility, all models were run with uniform inference parameters: temperature=0.0, top_p=1.0, frequency_penalty=0.0, and presence_penalty=0.0. By strictly controlling these settings, we enabled a direct and equitable comparison of each model’s extraction performance. Example code for the prompt and parameter configuration is provided in Figure 1.

**Figure 1.**
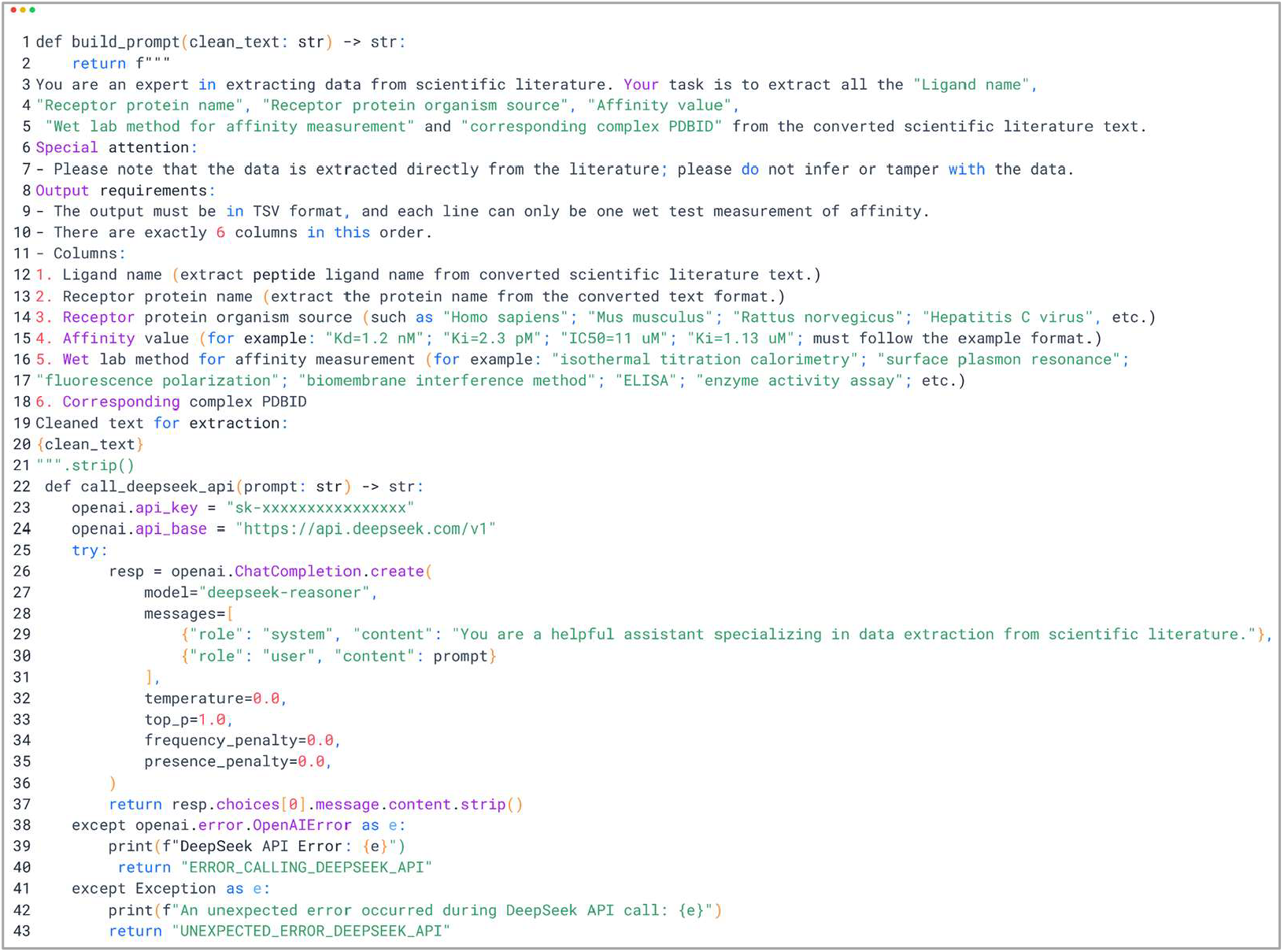
Prompt for extracting affinity data and model parameter settings (using Deepseek-reasoner as an example)

This evaluation workflow enabled us to systematically assess the performance differences among various LLMs when processing complex scientific texts, thereby providing critical insights into their suitability for large-scale, automated mining of protein–peptide interaction data. The evaluation results also offer guidance for PPIKB’s future LLM selection and prompt-engineering optimization, supporting its ongoing data expansion and iterative updates.

### 3. System Architecture and Workflow of PPIKB

The PPIKB platform is built upon a modern open-source technology stack and deployed on a Linux operating system. Its backend is powered by a high-performance Nginx web server, a MySQL database (version 5.5.29), and PHP, ensuring robust performance, security, and scalability. The web frontend is implemented using HTML, PHP, CSS, and JavaScript, providing users with an intuitive, interactive environment for data querying, visualization, and analysis.

At the core of PPIKB lies the data extracted from research articles and patents. These raw data are first processed by large language models (LLMs) for initial extraction, then subjected to rigorous manual validation and curation before being loaded into the MySQL database. Through its web interface, PPIKB offers multiple visualization modes (e.g., textual summaries, crystal structures, images) and various browsing options (including research article entries, patent entries, and structural entries, with category-based navigation). Users can explore the database via a suite of search tools— Text Search, Structure Search, and Quick Search—and leverage integrated sequence- and structure-based similarity search engines and editing tools, such as Foldseek^37^, BLAST^38^, SMILES Similarity, and the Ketcher Structure Editor. These capabilities enable rapid, sensitive similarity searches and comparative analyses based on peptide sequence or structural features, facilitating the identification of related peptides and protein targets. A schematic overview of the PPIKB construction workflow and detailed system architecture is presented in Figure 2.

**Figure 2.**
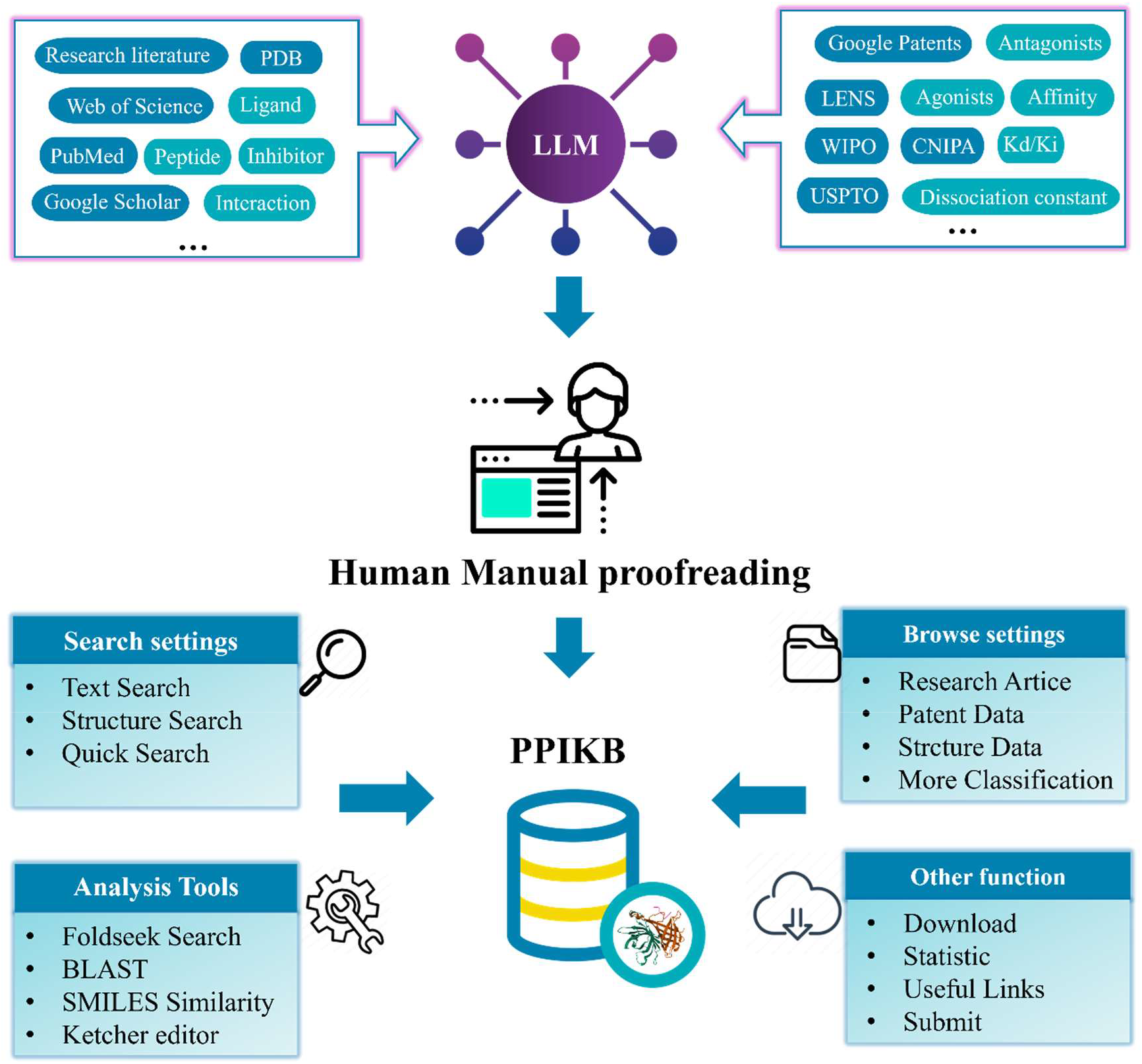
A brief overview of the construction process and system architecture of PPIKB

## Results

### 1. Comparison with existing databases

To highlight the advantages of PPIKB in terms of data coverage and content uniqueness, this study systematically compared it with several well-known protein-peptide/protein interaction databases (pepBDB^34^, PDBbind^35^, PepBind^32^ and Propedia V2.3^33^) in terms of the following key indicators: total entries, analysis and search tools, affinity data entries, structure information and PDB ID. The comparison results are summarized in Table 1.

**Table 1.**
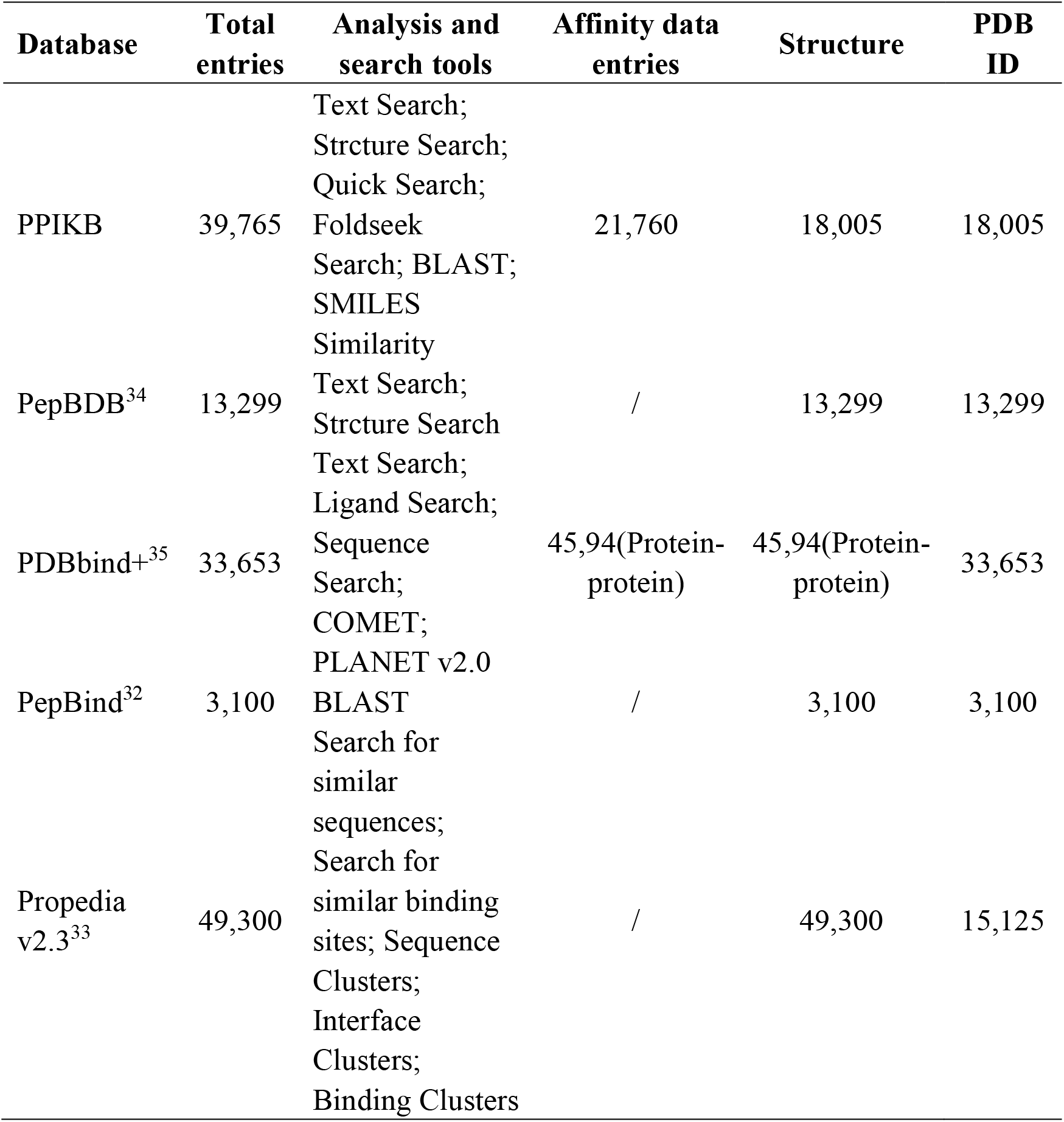
Comparison of PPIKB with other databases.

In addition, we also compared the number of protein-peptide interaction related data entries among PPIKB, pepBDB^34^ and PDBbind+^35^. As shown in Figure 3, the total number of entries and the overlap between the three are as follows: PPIKB contains a total of 39,765 protein-protein interaction records; pepBDB contains a total of 13,299 entries (structure only, no affinity)^34^, and all of them have been included in PPIKB; PDBbind+ contains 4,594 entries^35^. In the intersection of the three, 1,341 records appear in all three databases; another 1,947 records overlap between PPIKB and PDBbind+. The results in Table 1 show that PPIKB not only covers a wider range of protein-peptide interaction data, but also contains a large number of unique entries that have not been found in other databases. Compared with pepBDB^34^, which mainly focuses on structural information, PPIKB achieves more comprehensive information coverage by integrating three-dimensional structure data and quantitative binding affinity data extracted from massive research articles and patents. This dual focus on structure and affinity data allows researchers to obtain both atomic-level structural elucidation and quantitative interaction measurements in a single repository. On the other hand, although PDBbind^35^ is a mature repository for protein-ligand complex affinity data, its coverage of peptide-specific entries is more limited compared to PPIKB. With richer data entries and a large number of experimentally verified affinity values, PPIKB has become a more comprehensive and valuable tool platform in the fields of protein-peptide interaction research, drug discovery, and protein engineering.

**Figure 3.**
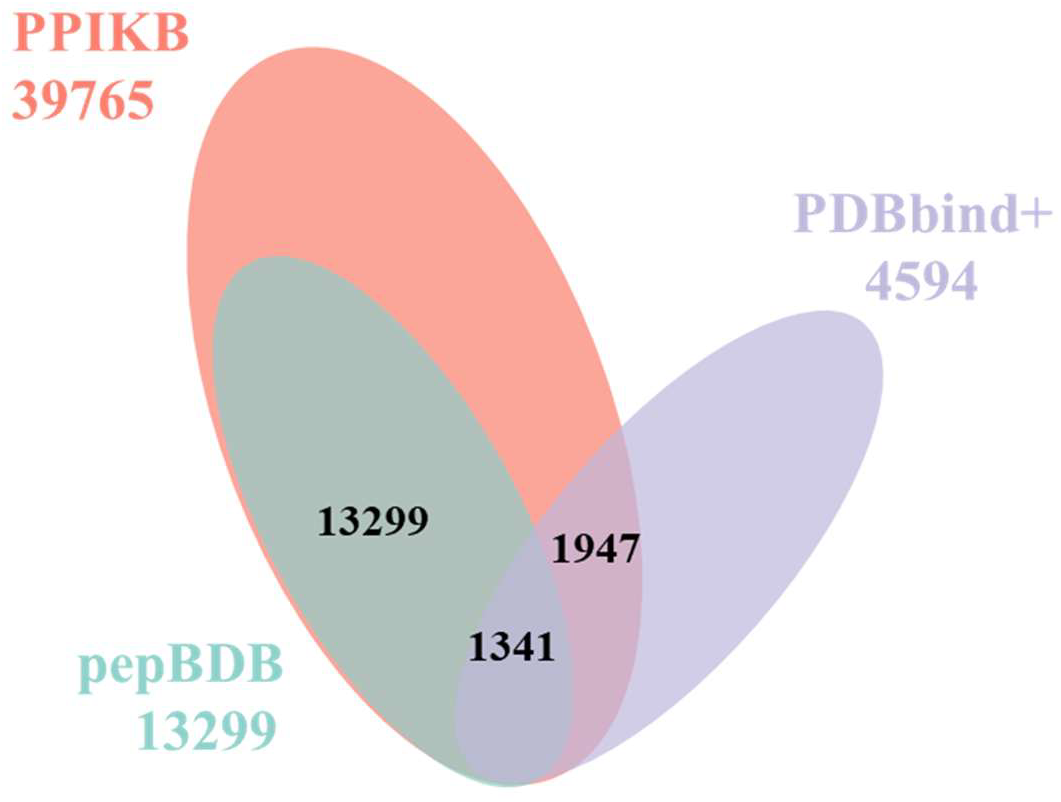
Venn diagram comparing protein-peptide/protein entries in PPIKB, pepBDB^34^ and PDBbind+^35^ databases

### 2. Detailed information entries in PPIKB

Detailed information in PPIKB is mainly divided into two categories. The first category is comprehensive information data on quantitative affinity between proteins and peptides, which is subdivided into five subcategories: target protein information (including protein name, protein sequence, organism source, subcellular localization, UniProt^42^ ID, and protein-protein interaction network information), basic peptide information (including peptide name, peptide sequence, SMILES, chemical modification, cyclization method, N-terminal modification, C-terminal modification, amino acid distribution), peptide physicochemical properties (molecular weight, aliphatic index, aromaticity, average rotatable bond, charge, isoelectric point, number of hydrogen bond acceptors, number of hydrogen bond donors, topological polar surface area, X_logP_energy), interaction information (affinity, affinity determination method, PDB_ID, type) and reference information (such as title, publication year, PMID, DOI, etc.). This type of affinity-related data mainly comes from published research articles and authorized patents, while structural information and protein sequences are compiled from the Protein Data Bank^31^ and UniProt databases^42^, respectively. The second type of information covers the protein-peptide structure comprehensive information data compiled from the PDB database^31^, including four types of information: protein information (such as protein chain name, protein sequence), peptide information (peptide sequence, chain name), interaction information (detailed contact details between proteins and peptides at the residue level, these contact information is calculated using the PLIP^40^ and reference information (such as title, PMID, DOI, etc.). Figures 4 and 5 show the visualization page of the comprehensive information of protein-peptide quantitative affinity and the visualization page of the comprehensive information of protein-peptide structure in Figure 4, respectively.

**Figure 4.**
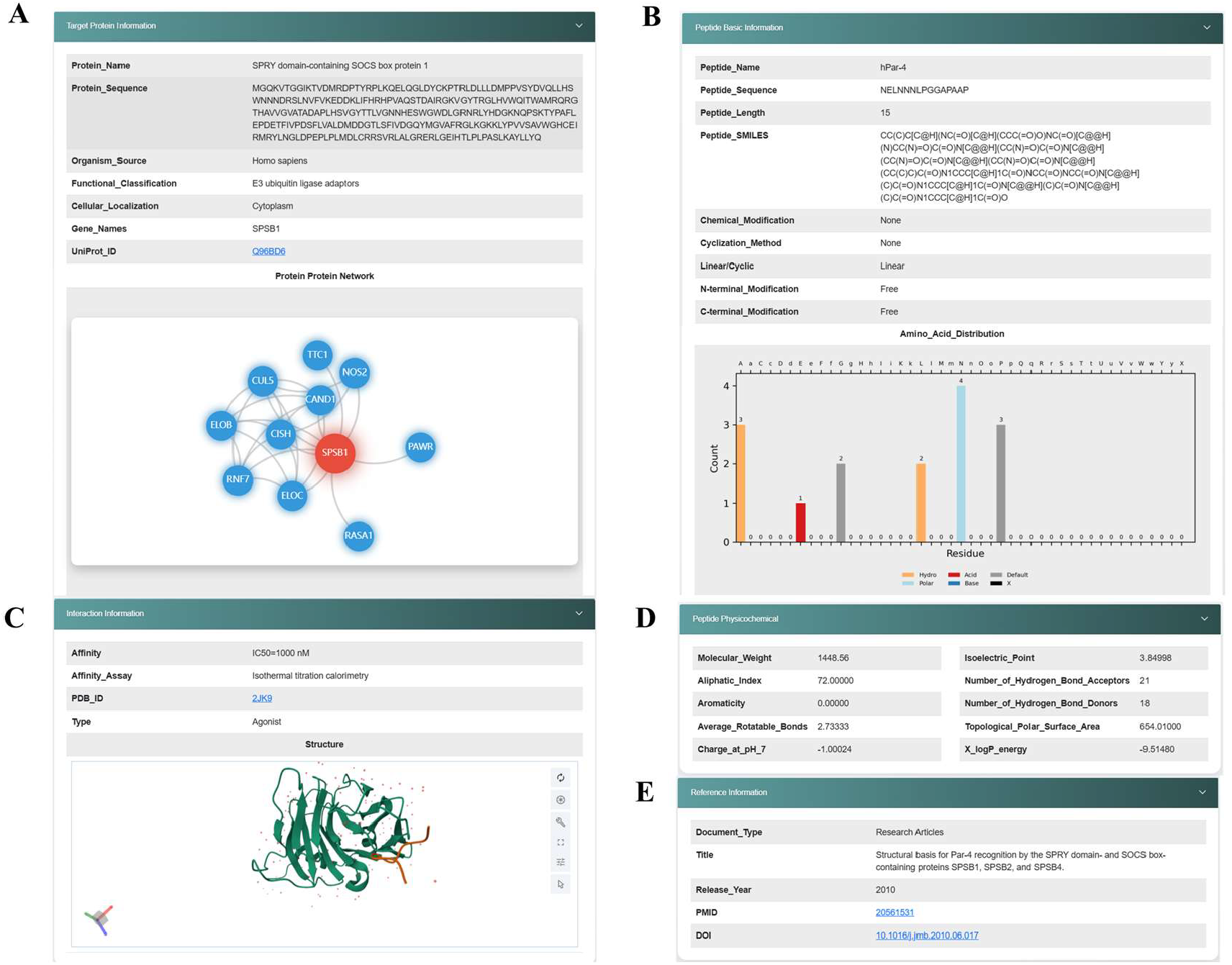
Visualization page of comprehensive information of protein-peptide quantitative affinity. **A**. Target protein information **B**. Basic peptide information **C**. Interaction information **D**. Peptide physicochemical properties **E**. Reference information

**Figure 5.**
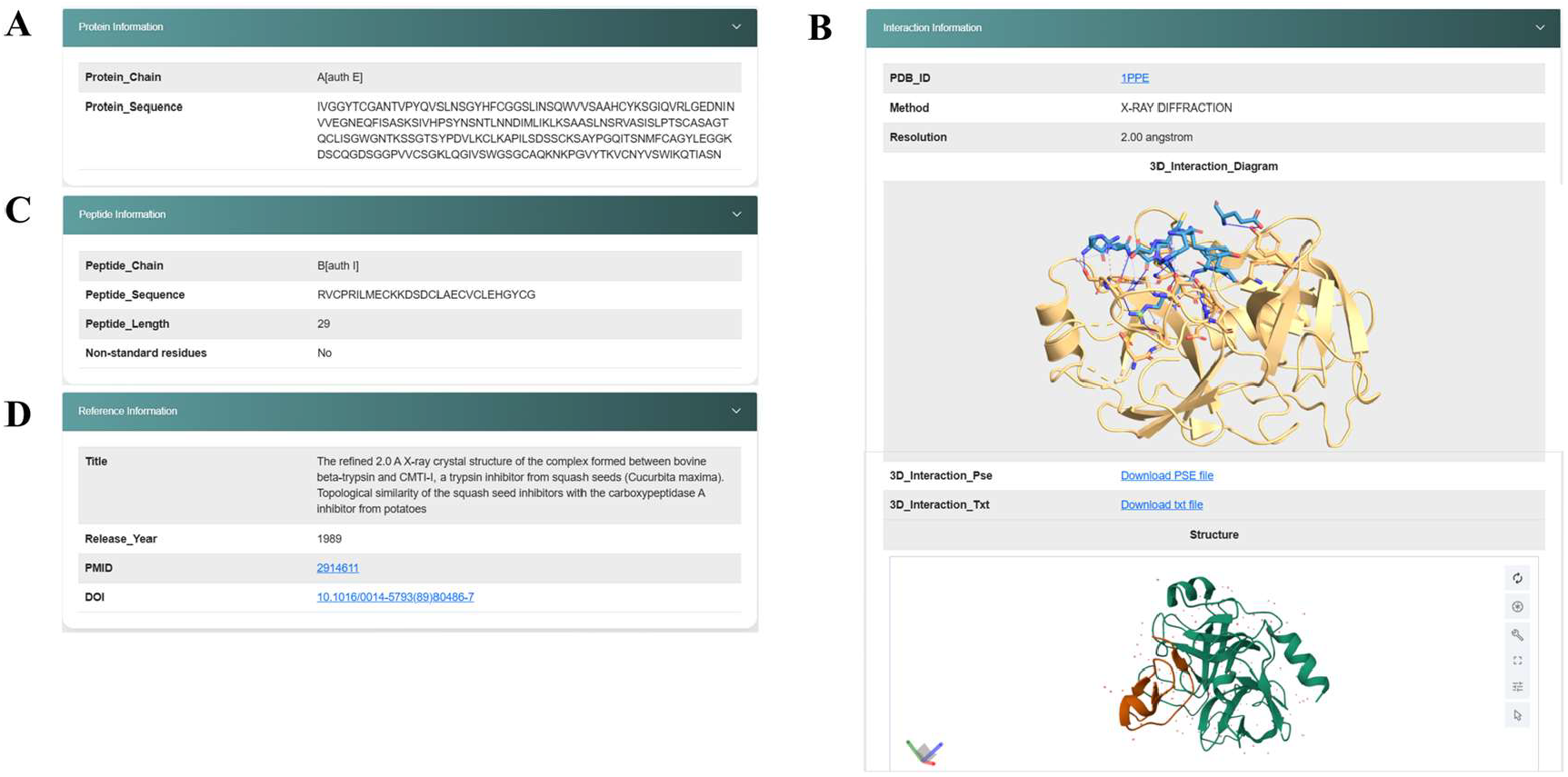
Visualization page of comprehensive information of protein-peptide structure. **A**. Protein information **B**. Interaction information **C**. Peptide information **D**. Reference information

### 3. Website Functions

To achieve efficient and customized search of PPIKB data, we have integrated dual search functions based on text and structure. In the Text Search module, users can perform precise queries on multiple fields, such as specifying affinity parameters (Kd, IC50, Ki) under Affinity Information and their specific units and numerical thresholds, thereby optimizing the search results, as shown in Figure 6. Structure Search module allow users to perform combined queries on crystallographic data, such as combining Method, Peptide Length Range, Residue Modification, Resolution Range, etc.

**Figure 6.**
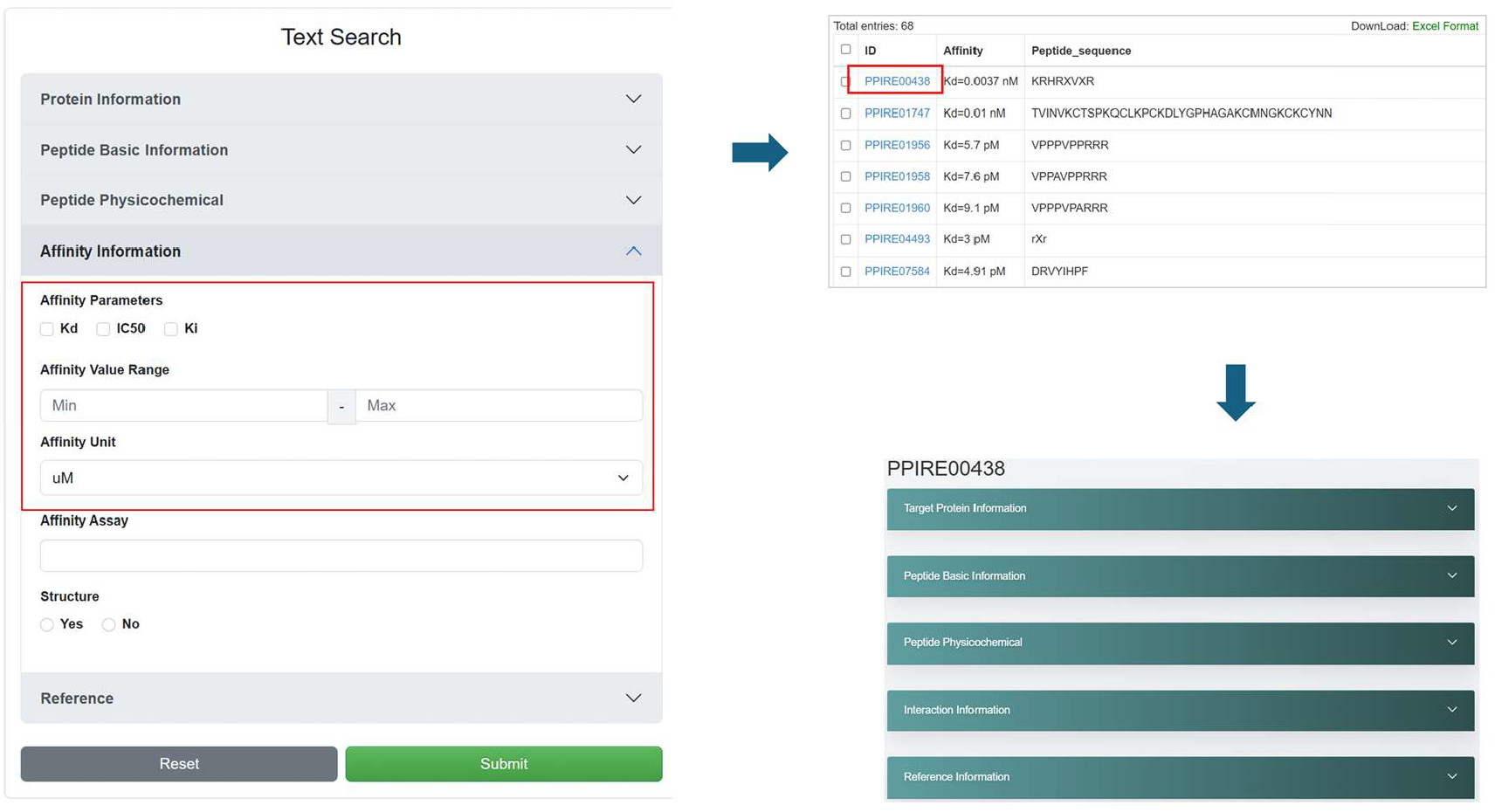
Example of using Text Search

In addition to these core search modes, PPIKB also integrates several widely used bioinformatics tools-FoldSeek^37^, BLAST^38^, SMILES-based similarity search, and Ketcher Structure Editor structure drawing to support users to conduct comprehensive and integrated analysis. For example, the FoldSeek^37^ interface allows users to perform a variety of structure-based and sequence-based search operations, and can directly link to the results of PPIKB, and can select appropriate output results according to parameters, as shown in Figure 7. For specific usage operations, see the help page (https://ppikb.duanlab.ac/static/help.php). To further facilitate the research community’s access to and contribution to our resources, we have also developed a dedicated data download (https://ppikb.duanlab.ac/downloads), submission page (https://ppikb.duanlab.ac/static/submit.php) and Useful Links page. The download page provides a convenient way to access all selected datasets with one click, while the submission page facilitates seamless data exchange and sharing between researchers and our team. Taken together, these carefully designed features together make PPIKB a flexible and interactive platform for exploring protein-peptide interaction data.

**Figure 7.**
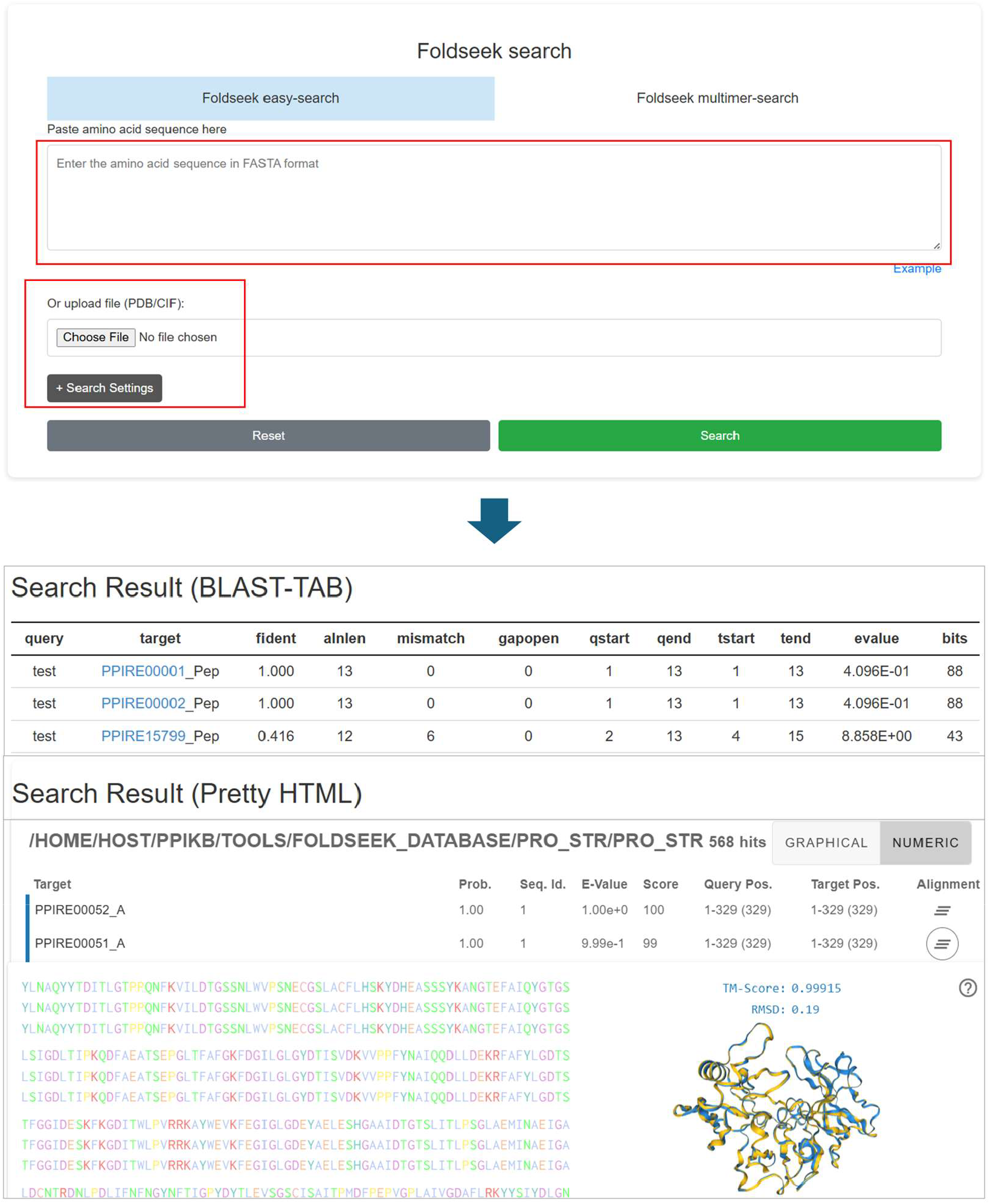
Example of using Foldseek

### 4. Evaluation of LLM data extraction accuracy

As shown in Figure 8, we conducted a comparative analysis of four large language models (Deep Seek-Reasoner, O3-Mini-2025-01-31, Gemini-1.5-Pro, and Claude-3-5-Haiku-20241022) on a classification task with 150 samples, and the results showed significant differences in accuracy among the models. DeepSeek-Reasoner performed best with an accuracy of 70% (105/150), followed by the lightweight O3-Mini-2025-01-31 with an accuracy of 66% (99/150). In contrast, the general-purpose Gemini-1.5-Pro and Claude-3-5-Haiku-20241022 achieved accuracies of 58% (87/150) and 36% (54/150), respectively. These findings highlight the potential advantages of retrieval-enhanced or domain-optimized architectures in structured classification tasks: Deep Seek-Reasoner’s superior performance may be partly due to its potential integration of high-fidelity structured information embeddings, which enhances the model’s ability to capture complex molecular relationships. In contrast, the lower accuracy of Claude-3-5-Haiku-20241022 reflects the limitations of general models that have not been fine-tuned for specific domains on this type of task, suggesting the need for targeted retrieval mechanisms or additional task-specific training. Notably, the small performance gap between Deep Seek-Reasoner and its lightweight counterpart suggests that computationally efficient models have the potential to approach the accuracy of larger and more complex systems. Future research should further explore integration strategies, advanced hint engineering techniques, and the trade-offs between model complexity, inference speed, and prediction reliability, especially in high-risk application scenarios such as drug development and protein engineering. In general, although the current LLM has not yet demonstrated extremely high accuracy in the direct extraction of protein-peptide affinity data and still requires a certain degree of manual verification, its processing efficiency (such as extraction speed, handling of multilingual patent documents, etc.) has been significantly improved compared to pure manual operations.

**Figure 8.**
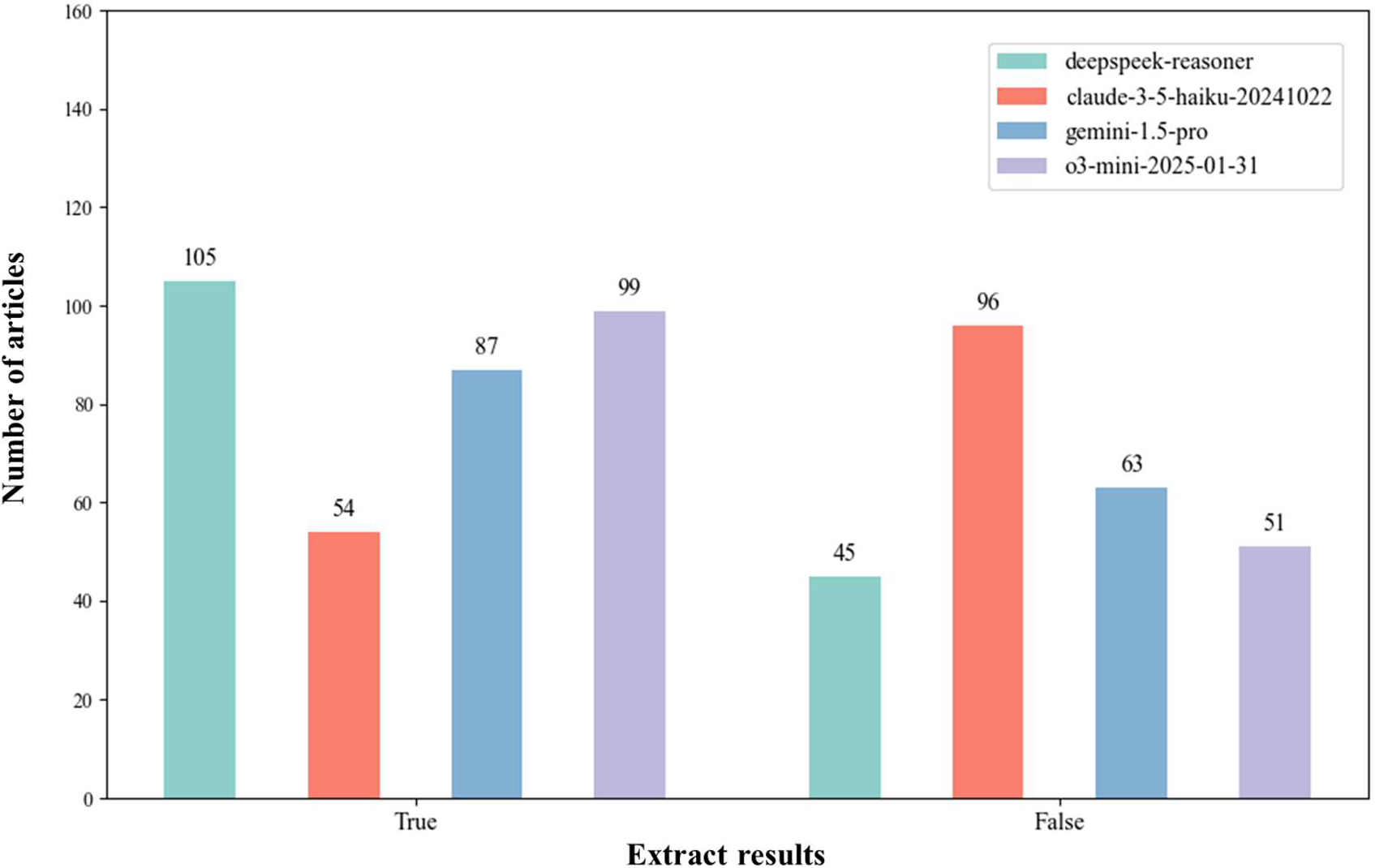
The results of data extraction by different LLM models. T means the data was extracted correctly, and F means the data was not extracted.

### 5. Data summary and statistics

The current version of PPIKB contains two types of data information. The first type is protein-peptide quantitative affinity comprehensive information data, with a total of 21,760 entries. The main affinity data comes from 483 patents and 4,241 research papers. As shown in Figure 9A, the publication years of research articles and patents in PPIKB span from 1,972 to 2,025, covering more than half a century of scientific research and innovation results. Although the number of early entries was relatively limited, it has steadily increased since the 1990s and reached a peak between 2,010 and 2,015, reflecting the significant increase in research enthusiasm and technological innovation in this field. This extensive and continuous time series distribution not only demonstrates the depth and breadth of PPIKB in data collection, but also provides a solid foundation for subsequent trend analysis, knowledge graph construction and discipline evolution research. So far, this study has included 2,013 unique protein targets, covering a variety of biological sources such as humans, mice, rats, viruses and bacteria. As shown in Figure 9B, among all the entries, targets from Homo sapiens accounted for the highest proportion, about 66%; followed by “Other organisms” (12%), viruses (7%), mice (6%), rats (5%) and bacteria (4%). Among all human targets, the top three protein targets in the top 30 are angiotensin-converting enzyme, dipeptidyl peptidase 4 and E3 ubiquitin-protein ligase Mdm2, all of which belong to enzymes; followed by Bcl-2-like protein 1, whose target type is regulatory proteins, as shown in Figure 9C. The second category is protein-peptide structure comprehensive information data, which contains a total of 18,005 entries, corresponding to 10,418 research papers. As shown in Figure 9D, structure determination mainly relies on X-ray diffraction (XRD), accounting for 89.83%; electron microscopy (EM) accounts for 4.83%; the remaining methods include solution nuclear magnetic resonance (solution NMR) 5.25%, solid-state NMR 0.04%, neutron diffraction 0.02%, powder diffraction 0.02% and electron crystallography 0.01%. Figure 9E compares the resolution distribution of the two technologies of XRD and EM: the resolution determined by XRD is mainly concentrated in the range of 1.5-2.5 Å, with a median of about 2.1 Å, and a compact distribution; while the EM resolution is mostly distributed in 2.5-3.5 Å, and has a significant extension to higher values, with a median of about 3.4 Å. Overall, the structural resolution obtained by XRD is significantly better than that of EM, providing important support for high-precision structural analysis. For more statistical analysis, see https://ppikb.duanlab.ac/static/statistic.php.

**Figure 9.**
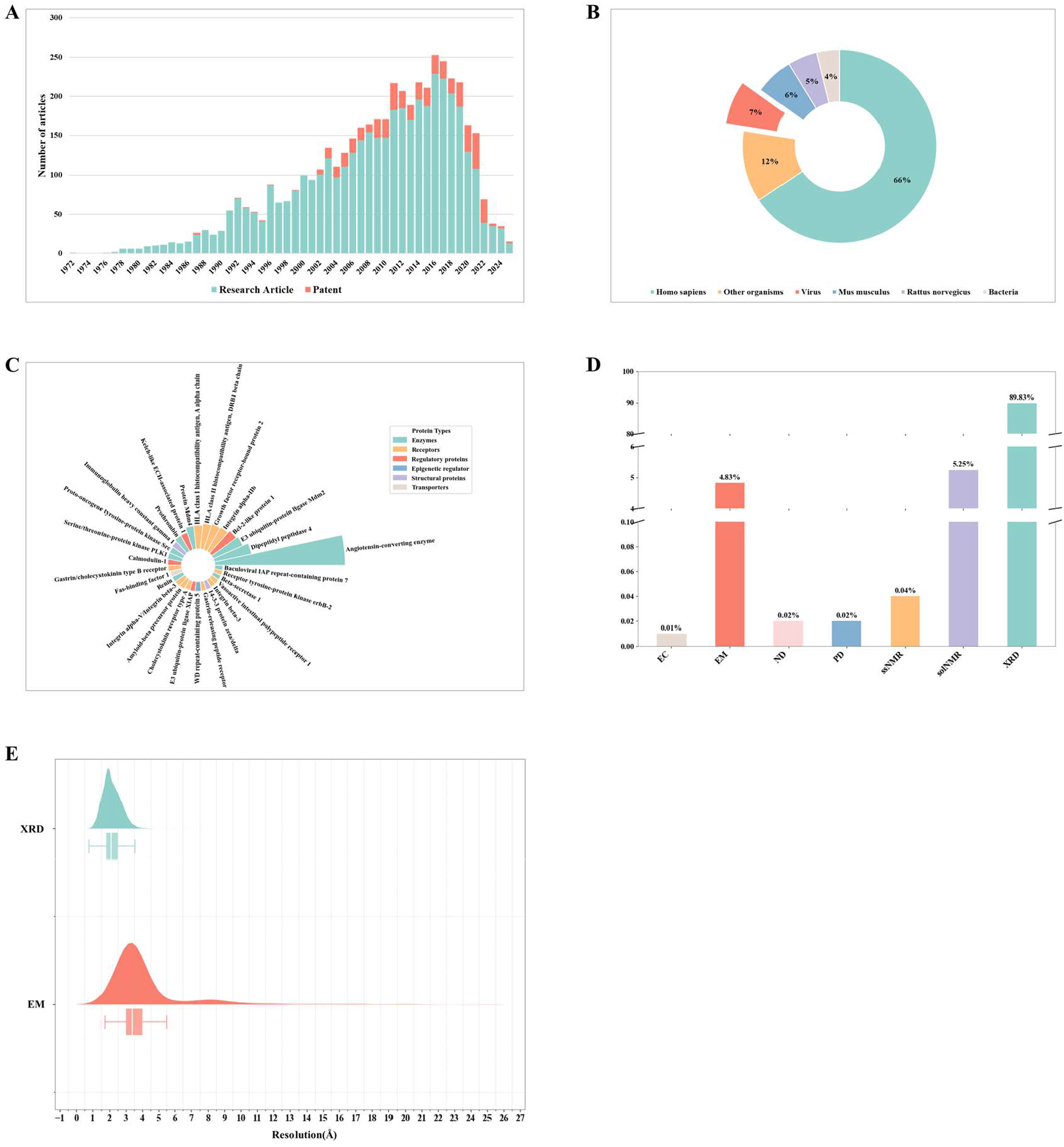
PPIKB data statistical analysis. **A**. Distribution of publication years of research articles and patents **B**. Distribution of species origins of target proteins. **C**. Distribution of the top 30 target proteins from Homo sapiens. **D**. Distribution of protein–peptide crystal structure determination methods. **E**. Distribution of resolutions for crystal structure determination by X-XRD and electron EM. **Abbreviations**: **EC**: electron crystallography; **EM**: electron microscopy; **ND**: neutron diffraction; **PD**: powder diffraction; **ssNMR**: solid-state nmr; **solNMR**: solution nmr; **XRD**: x-ray diffraction

## Conclusion

As the first comprehensive protein-peptide interaction knowledge base built based on a combination of large language models and manual verification, PPIKB fills the gap in the systematic integration of peptide binding affinity data and structural data in existing resources. By incorporating data from a total of 4,241 literature and patents from 1972 to this year, PPIKB ultimately provides 20,234 manually verified quantitative affinity entries and integrates 18,005 protein-peptide complex structure information, covering 2,013 protein targets from various biological sources such as humans, mice, rats, viruses and bacteria. Compared with existing databases that only focus on structural information, PPIKB achieves dual integration of affinity and structural data, allowing researchers to simultaneously obtain atomic-level structural analysis and quantitative interaction measurement results on the same platform; compared with traditional protein-ligand databases, PPIKB focuses on peptide-protein interactions, filling the data gap in this field.

In terms of technical routes, we first introduced a large language model for large-scale literature and patent mining, combined with strict manual verification, which significantly improved the efficiency and quality of data extraction. Although various LLMs (such as DeepSeek, Chatgpt, Gemini, and Claude) perform differently when extracting affinity data, they are all superior to pure manual operations in terms of processing speed and coverage. In terms of system architecture, PPIKB is based on a mature open source technology stack, achieving high performance, high scalability, and good user experience. The platform has built-in analysis tools such as Foldseek, BLAST, and SMILES Similarity, which support users to conduct multi-dimensional search and data mining, providing strong support for peptide drug development, protein engineering, and systems biology research.

## Future update plan

In order to continue to promote the development of PPIKB, we will take a number of measures in future updates: first, run an automatic update program every three months, and the update log will be displayed at https://ppikb.duanlab.ac/update.php to timely integrate the latest protein-peptide interaction research results; second, make special fine-tuning of the large language model to further improve the ability to accurately extract data from the literature; in addition, protein-protein affinity data will be introduced to enrich the database content. We sincerely look forward to the valuable suggestions of researchers to jointly promote the in-depth development of peptide drug development, protein engineering and systems biology research.

## Data availability

https://ppikb.duanlab.ac/

## Acknowledgment

This project was supported by the Macao Science and Technology Development Fund (Grant No. 0151/2024/RIA2) and the internal grant from Macao Polytechnic University (RP/FCA-07/2024).

## Funder Information Declared

Macao Science and Technology Development Fund, No. 0151/2024/RIA2 internal grant from Macao Polytechnic University, RP/FCA-07/2024

